# At-home blood collection and stabilization in high temperature climates using *home*RNA

**DOI:** 10.1101/2022.03.23.485526

**Authors:** Lauren G. Brown, Amanda J. Haack, Dakota S. Kennedy, Karen N. Adams, Jennifer E. Stolarczuk, Meg G. Takezawa, Erwin Berthier, Sanitta Thongpang, Fang Yun Lim, Damien Chaussabel, Mathieu Garand, Ashleigh B. Theberge

## Abstract

Expanding whole blood sample collection for transcriptome analysis beyond traditional phlebotomy clinics will open new frontiers for remote immune research and telemedicine. Determining the stability of RNA in blood samples exposed to high ambient temperatures (>30°C) is necessary for deploying home-sampling in settings with elevated temperatures (e.g., studying physiological response to natural disasters that occur in warm locations or in the summer). Recently, we have developed *home*RNA, a technology that allows for self-blood sampling and RNA stabilization remotely. *home*RNA consists of a lancet-based blood collection device, the Tasso-SST™ which collects up to 0.5 mL of blood from the upper arm, and a custom-built stabilization transfer tube containing RNA*later™*. In this study, we investigated the robustness of our *home*RNA kit in high temperature settings via two small pilot studies in Doha, Qatar (no. participants = 8), and the Western and South Central USA during the summer of 2021, which included a heatwave of unusually high temperatures in some locations (no. participants = 11). Samples collected from participants in Doha were subjected to rapid external temperature fluctuations from being moved to and from air-conditioned areas and extreme heat environments (up to 41°C external temperature during brief temperature spikes). In the USA pilot study, regions varied in outdoor temperature highs (between 25°C and 43.4°C). All samples that returned a RNA integrity number (RIN) value from the Doha, Qatar group had a RIN ≥7.0, a typical integrity threshold for downstream transcriptomics analysis. RIN values for the Western and South Central USA samples (n=12 samples) ranged from 6.9-8.7 with 9 out of 12 samples reporting RINs ≥7.0. Overall, our pilot data suggest that *home*RNA can be used in some regions that experience elevated temperatures, opening up new geographical frontiers in disseminated transcriptome analysis for applications critical to telemedicine, global health, and expanded clinical research. Further studies, including our ongoing work in Qatar, USA, and Thailand, will continue to test the robustness of *home*RNA.

## INTRODUCTION

Blood is a useful biofluid for transcriptome analysis, as it is relatively non-invasive to obtain and provides ample information for biomarker identification and analysis 1. Consequently, whole blood RNA transcript analysis has emerged as an avenue for realizing the goals of personalized medicine ^2,3^. Key biomarkers discovered through blood transcriptomic studies have potential applications in novel therapeutics and diagnostics ^4–8^ and have provided new avenues for disease monitoring of chronic ^9,10^, autoimmune ^11,12^, and infectious diseases ^13^. By combining home-sampling technologies and transcriptomics, one can further expand upon these applications, opening up both biomarker discovery through discovery-based research studies, and bring applications such as gene expression-based disease monitoring and diagnostics to home-based tests. Further, a technology that allows for remote blood transcriptomics would be enabling for increasing access to rural, remote, or underserved communities, for both research studies and telemedicine, and could be a powerful technology for enhancing global health efforts^2^.

Conducting remote blood transcriptomics research presents several challenges. For example, while blood is relatively non-invasive to collect, conventional blood collection is limited to phlebotomy. Logistically, relying on phlebotomy for collection geographically confines blood collections to clinic/lab locations or research institutes. Several self-blood collection technologies have been developed for remote use, including fingersticks 8, dried blood spots ^14–16^, and lancet-based technologies that collect blood from the upper arm ^17–21^, such as the Tasso-SST™ device that we use in our home-sampling and RNA stabilization kits (including the work in this manuscript). Another major challenge of remote blood transcriptomics is stabilization of whole blood RNA to minimize RNA degradation due to spontaneous degradation via auto hydrolysis or degradation due to the presence of ribonucleases ^22^. Toma *et al*. developed an at-home blood collection kit that collects 50 μL of capillary blood via fingerstick and mixes with a proprietary RNA preservative solution at a 1:4 ratio ^8^. To test the ability of the RNA preservative solution to preserve blood transcriptome, Toma *et al*. collected blood samples from participants (n=3) and stored at ambient temperature (22°C) or -80°C for 0, 7, 14, and 28 days, with and without shipping. They found high correlation coefficients between gene expression levels from the samples in all conditions, demonstrating the efficacy of the RNA stabilization solution to preserve RNA integrity over 28 days^8^.

To address this burgeoning need and develop technology for higher blood volumes than are possible with fingersticks, we have recently developed *home*RNA, a kit for the self-collection and stabilization of whole blood RNA, for use in disseminated whole blood transcriptome applications ^23^. *home*RNA uses a blood collection device, the Tasso-SST™, that collects capillary blood from the upper arm via a lancet; the blood collection tube from the Tasso system then interfaces with a custom engineered stabilizer tube that holds liquid stabilizer solution (RNA*later™*) ^23^. Previously, we demonstrated successful stabilization of RNA extracted from samples collected with *home*RNA in 47 participants in 10 USA States in a pilot feasibility study for validation of *home*RNA for remote blood collection and RNA stabilization. However, most of these samples were collected in regions with relatively moderate summer temperatures or during the fall/winter and thus we sought to test the *home*RNA sampling kit in regions with elevated temperatures. Our present study builds on a small body of prior work examining the effects of blood storage temperature on RNA quality; prior work largely focused on the effects of cold storage and did not examine temperatures above 25°C ^24–26^.

Establishing means to remotely probe the human whole blood transcriptome in elevated temperature settings has important implications, including enabling clinical research or personalized medicine applications in areas with high temperature and studying diseases/events that are specific to warm climates or warm times of year. For example, regions more highly burdened by tropical diseases would benefit from a better understanding of the stability of RNA because many of these areas often have persistently elevated temperatures. Further, even in regions with more moderate climates for part of the year, researchers may wish to conduct remote sampling studies in the summer months, particularly when tied to a specific seasonal event. For example, a study aiming to understand wildfire smoke exposure in the Western and South Central USA would necessitate sampling in the summer months where temperatures are elevated, as these months are when wildfires occur. Overall, the ability to reliably stabilize RNA with *home*RNA in high temperature settings will open up remote transcriptome studies to regions with a warmer climate, whether seasonally or permanently. We present here two small pilot studies exploring the application of *home*RNA in high temperature settings. In the first pilot study, *home*RNA was used by a small group in the city of Doha, Qatar before the sample was exposed to simulated shipping conditions of fluctuating temperatures. In the second study, *home*RNA was used at a regional scale, where participants used *home*RNA in their own homes during the summer months in several Western and South Central USA states, which included regions that are typically very hot (e.g., Reno, Nevada and Southern California) as well as the Pacific Northwest during the June 2021 heat wave. For samples collected from both studies, we examine quality control metrics typical to determining suitability of isolated RNA for downstream RNA transcript analysis, including the RNA integrity number (RIN) and total cellular RNA yield. In both the small pilot study in Qatar, and the small pilot study in the Western and South Central USA, we found satisfactory RIN values (RIN ≥7) in the majority of samples (84%, 16 out of 19 samples), with a RIN of 6.9 in the remaining 3 samples. Overall, our data demonstrate successful RNA stabilization as a function of yield and RIN values using *home*RNA in high temperature settings.

## MATERIALS AND METHODS

### *home*RNA for the collection and stabilization of whole blood cellular RNA

#### Design and assembly of homeRNA

The fabrication of the RNA stabilizer tube as an interface to the Tasso-SST™ blood tube has been described previously ^23^. Briefly, the RNA stabilizer tube was designed to hold the RNA*later*™ stabilization solution and connect to the Tasso-SST™ blood collection tube for remote self-blood collection and RNA stabilization. The Tasso-SST™ blood collection tube does not contain an anticoagulant. We previously showed that Tasso-SST™-mediated capillary blood collection, despite not having an anticoagulant, did not result in major compromise of RNA quality as measured by RIN values (23). At the time of the development of the *home*RNA kit, the Tasso-SST™ was the only commercially available Tasso product for collecting liquid samples; we note that we do not use the serum separator tube (SST) feature. The stabilizer tube was injection molded out of polycarbonate (PC: Makrolon 2407) by Protolabs, Inc (Maple Plain, MN). Details of the stabilizer tube components and design files can be found in Haack, Lim *et al*. ^23^. Prior to assembly of the stabilizer tube, all components were first cleaned via sonication in 70% ethanol (v/v) for 30 min and air dried. The stabilizer tube was filled with 1.3 mL RNA*later*™ (Thermo Fisher) stabilization solution and capped. The stabilizer tube insert was designed to hold the stabilizer tube containing blood in a 50 mL conical tube during transport. The insert also allows for facile centrifugation of the sample in the 50 mL standard conical tube when it arrives in a laboratory setting, as it centers and immobilizes the *home*RNA sample tube relative to the 50 mL tube. The insert was injection molded out of polycarbonate (PC: Makrolon 2407) by Protolabs, Inc (Maple Plain, MN). The Instructions for Use (IFU) is included in the SI of this manuscript for easy reference (Appendix 1) and is an adaption of the previously reported IFU in the supporting information of Haack, Lim *et al*. ^23^. All kit components were placed in a rigid custom design mailer box fabricated via die-cutting (The BoxMaker, Inc.). ^23^

### High temperature fluctuations in *home*RNA collected samples: a small pilot study in Doha, Qatar

#### Participant recruitment and demographics

Healthy adult volunteers (18 years or older) were recruited in Doha, Qatar via word of mouth under a protocol approved by Sidra Medicine Institutional Review Board study number 1609004823. Written informed consent was obtained from all participants. In total, the study enrolled 8 participants between the ages of 29 and 53 (7 male and 1 female).

#### Temperature fluctuations of homeRNA collected and stabilized blood samples

Participants were asked to collect and stabilize 1-2 whole blood samples using *home*RNA in a laboratory setting. The stabilizer vial containing the stabilized blood was placed in the 50 mL conical tube with the insert provided in the *home*RNA kits (see *design and assembly of homeRNA* above). Next the 50 mL tube containing the stabilized blood samples was placed in a sealed bag (provided in the kit) inside the *home*RNA cardboard package, and then incubated at ambient temperature (21°C) for 27 hours to simulate the amount of time that might pass between blood collection at home and a scheduled pickup time from a courier service. Samples 1a (from participant 1 who was sampled twice) and 2 were frozen at -80°C after incubation at ambient room temperature for 27 hours. The remaining 6 samples, after 27 hours incubation at ambient temperature, underwent 3 hours of exposure to cycling temperatures in air-conditioned, indoor, and outdoor environments to simulate shipping conditions of the *home*RNA kit from remote site to lab. Each sample was exposed to a low of 21°C and a high of 41°C (with the elevated temperatures occurring as brief temperature spikes, see Table S1 and rationale for this temperature exposure/study design provided in the Results and Discussion) over the course of 27 -30 hours after blood collection and stabilization.

### *home*RNA collection during summer months: a small pilot study in the Western and South Central USA

#### Participant recruitment and demographics

Healthy adult volunteers (18 years or older) were recruited from US states that historically experience wildfire smoke (i.e., the Western USA) via word of mouth and digital ads under a protocol approved by University of Washington Institutional Review Board (STUDY00012463). 11 participants between the ages of 20 and 56 years old were enrolled (4 male and 7 female) and collected samples used in this study. Note: these samples were collected as part of a larger ongoing study our research group is performing to investigate the effects of wildfire smoke exposure on inflammation. These 12 samples (from 11 participants) were chosen to capture a range of temperatures including the warmest temperatures experienced by our participants in summer 2021. Written informed consent was obtained from all participants. Participants were located in 7 states, including California, Nevada, New Mexico, Texas, Oregon, Washington, and Wyoming. Samples were collected between June and July of 2021, including samples (n = 2) that were collected during the June 2021 heat wave in the Pacific Northwest.

#### Participant blood self-collection and stabilization

Participants were asked to self-collect and stabilize blood from their upper arm using the *home*RNA blood collection and stabilization kit. Each participant was also asked to complete a survey with each collection that included information about the ambient temperature during collection and the approximate blood level (see Figure S6). Participants were asked to package their stabilized blood samples to be returned to the lab for analysis using the provided return mailer bag (per instructions including containment in a 50 mL conical tube, plastic bag, and the *home*RNA kit cardboard box). Participants left their sample outside for pickup between the hours of 12:00 PM and 3:00 PM. All stabilized blood samples were mailed using next day delivery courier services (United Parcel Service, UPS). Returned samples were delivered to a -20ºC freezer and subsequently stored at -80ºC prior to RNA extraction.

#### Temperature data collection

Participants were provided with a temperature monitor (ThermPro) and were asked to record the temperature at the time of sampling to obtain the indoor temperature data (Fig. 3Bi). We collected the maximum temperature from the day the sample was picked up as the outdoor high on pickup day (Fig. 3Bii). Temperature data was obtained from the daily high reported from the closest weather station (as determined by the participant’s zipcode) on the day of the pickup. The weather station data was obtained from Weather Underground.

### RNA isolation and assessment of RNA integrity from *home*RNA-stabilized whole blood

#### RNA isolation

For both small pilot studies in Doha, Qatar and the Western and South Central USA, total cellular RNA was isolated using the Ribopure™ - Blood RNA Isolation Kit (Thermo Fisher) according to manufacturer’s protocol and eluted in two 50 μL aliquots. Prior to removing the stabilized blood from the collection tube, the 50 mL conical tube holding the *home*RNA device was briefly centrifuged 10 s at 50 g. Isolated RNA was stored at -80°C until ready for further analysis.

#### Assessment of yield and RNA integrity of total cellular RNA

For the pilot study in the USA, RIN values of the first 50 μL elution were obtained on a Bioanalyzer 2100 (Agilent). The Agilent 2100 Bioanalyzer uses proprietary, built-in software to calculate RIN values, which is based on the relative peak height and shape of the 18S and 28S rRNA fragments in the resulting electropherogram after separation. All samples were assessed using the RNA 6000 Nano Kits. Samples with returned concentrations below the quantitative range (<25 ng/µL) of the RNA 6000 Nano Kit (samples 1-8) were further analyzed using the RNA 6000 Pico Kit. For these low concentration samples, only values provided by the Pico Kit were used in the data analysis. RNA concentration was measured for both isolated RNA aliquots using a Cytation 5 Multi-Mode Reader (Agilent Biotek Instruments) with a Take3 Micro-Volume Plate at the wavelengths of 260, 280, and 320 nm. For the pilot study in Doha, Qatar, RNA integrity number (RIN) values and RNA concentration of the first 50 μL elution were obtained on a Bioanalyzer 2100 (Agilent) using the RNA 6000 Nano Kit (Agilent) for all samples. For samples analyzed in Doha, additional assays on the Pico Kit were not performed on samples with concentrations below the qualitative range of the Nano Kit.

## RESULTS AND DISCUSSION

### home*RNA at high temperatures: an overview of two pilot studies*

In this work, we tested the robustness of the *home*RNA kit by evaluating the stability of RNA transcripts in blood collected using the *home*RNA kits during high temperatures in the summer months (July-August) in two different regions: Doha, Qatar and the Western and South Central USA. In the Doha study, samples were collected in-lab with *home*RNA, followed by in-field sample exposure in a hot car to simulate conditions in a courier service.

The Western and South Central USA study employed remote collection with *home*RNA by participants in their home, and samples were then shipped back to the lab. A typical logistical process of collecting, stabilizing, shipping, and processing *home*RNA blood samples is outlined in Fig. 1. In each case, our study design was chosen based on how at-home clinical research studies would be performed in that region (i.e., a 3-hour car ride to simulate a courier service in Doha and overnight shipping via UPS in the USA). Specifically, in the Qatar pilot study we were interested in understanding if stabilized blood samples exposed to high temperature fluctuations in a region of the world that regularly experiences high temperatures would affect the quality of isolated blood RNA. The Qatar samples were collected in a laboratory setting which was air conditioned (to model the air-conditioned home environment typical in Qatar) and deliberately stressed in different temperature environments (i.e., collection at air-conditioned ambient temperature, exposure to external outdoor high temperatures, transportation conditions, etc. (Table S1)) to simulate shipping conditions that typically occur between remote blood collection and shipment back to the lab for analysis in Qatar. In Qatar, air conditioning is prevalent, and therefore samples may go from air-conditioned areas to extreme heat during shipping. Further, in Doha courier services (such as Uber) are a common method of moving packages around the city. The conditions we employed (Table S1) were chosen to mimic a typical car ride that the samples would experience in a courier service for shipments within the city itself; the length of the car ride (3 hours) is shorter than the time required to ship packages in the USA with UPS but is relevant for courier services in Doha. Subsequently, the Western and South Central USA pilot study experienced real-world shipping conditions in which participants were mailed *home*RNA kits from Seattle WA to various states in the Western region of the USA during the summer (Table 1), yielding data from participants in a range of regions and temperatures to further test the applicability of *home*RNA in multiple high temperature environments.

**Table 1:** Participant and sampling information for the Western and South Central USA pilot study.

| Participant | Self-Reported Blood Volume (μL) | Indoor Temperature at Sampling (°C) | Outdoor Temperature High on Pickup Day (°C) | Location of Sampling | Month of Sampling | Total Yield (ug) | RIN |
| --- | --- | --- | --- | --- | --- | --- | --- |
| 1A | 200 | 25.0 | 42.3 | Albany, OR | Late June | 0.7 | 6.9 |
| 1B | 300 | 26.3 | 35.6 | Albany, OR | Early July | 0.8 | 6.9 |
| 2 | 400 | 25.8 | 36.8 | Davis, CA | Late July | 1.1 | 7.1 |
| 3 | 300 | 23.2 | 28.7 | Seattle, WA | Early August | 0.8 | 7.4 |
| 4 | 300 | 24.0 | 33.4 | Dallas, TX | Early July | 0.7 | 7.3 |
| 5 | 200 | 23.1 | 35.3 | Laramie, WY | Late July | 0.5 | 7.6 |
| 6 | 200 | 24.5 | 34.8 | Pasadena CA | Mid July | 0.8 | 6.9 |
| 7 | 300 | 26.9 | 43.4 | Portland, OR | Late June | 0.5 | 7.4 |
| 8 | 400 | 27.2 | 32.3 | Reno, NV | Early August | 1.0 | 7.3 |
| 9 | 300 | 21.2 | 25.0 | Albuquerque, NM | Late June | 1.9 | 8.2 |
| 10 | 400 | 25.0 | 35.4 | Portland, OR | Late July | 1.8 | 7.6 |
| 11 | 400 | 23.0 | 25.5 | Portland, OR | Late July | 1.6 | 8.7 |

**Figure 1.**
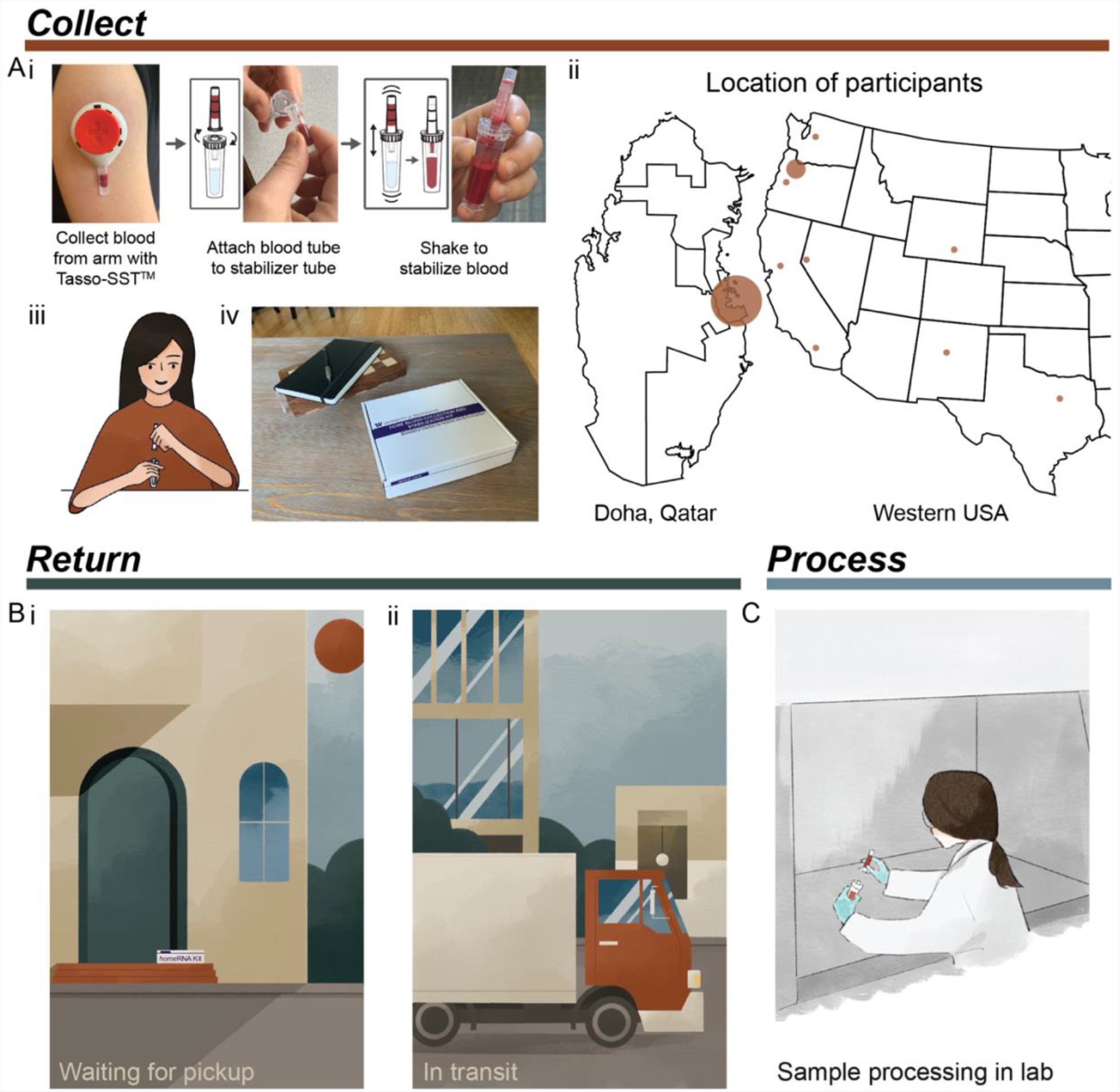
Typical process for using *home*RNA from collection to processing of samples. A) Collection and stabilization of blood using homeRNA. i) Process of collecting blood from the upper arm with Tasso device and stabilizing the sample with the *home*RNA custom stabilizer tube. Image was reprinted with permission from Haack, Lim *et al. home*RNA: A Self-Sampling Kit for the Collection of Peripheral Blood and Stabilization of RNA. *Anal. Chem*. 2021, 93, 39, 13196–13203. Copyright 2021 American Chemical Society.^23^ ii) Map depicting locations of participants in the small pilot studies conducted in Qatar and the Western and South Central USA. Note, participants in Qatar collected samples themselves in a lab setting. iii) illustration demonstrating a participant connecting the Tasso blood tube and the *home*RNA stabilizer tube, iv) *home*RNA kit on a coffee table, depicting a home setting for blood collection. B) Illustration demonstrating two possible locations where samples may be exposed to high temperatures including i) located on a front porch waiting to be picked up (which was the pickup location for many participants in the Western and South Central USA pilot study) and ii) in transit in a delivery truck or courier service. C) Final step of sample processing for downstream analysis in a laboratory setting.

The robustness of the *home*RNA kit to elevated ambient temperature exposures was validated by assessing isolated blood RNA quality with RNA integrity number (RIN) values. RIN values range between 1 and 10, with 1 referring to a completely degraded sample and 10 referring to an intact sample ^27^. RIN values ≥7.0 are standard QC cut-offs for genome wide transcriptional profiling ^28^. However, useful RNA sequencing data can still be obtained from more degraded samples (RIN values between 4 and 7) ^29^.

### home*RNA for high temperature regions: a pilot study in Doha, Qatar*

To understand the robustness of the *home*RNA device for use in clinical studies performed in Doha, Qatar, 8 participants in Doha collected and stabilized their own blood in late August 2021. After blood collection and stabilization with the *home*RNA kit, each sample was incubated at room temperature (21°C) for 27 hours, simulating time between remote blood collection and next-day sample pick up for shipment. After sitting out at room temperature, the stabilized blood samples were then exposed to fluctuating external temperatures in air-conditioned, indoor, and outdoor environments, as outlined in Fig. 2A and Table S1, by driving the samples around Doha in a car. This setup mimics the change in environment and temperatures that the stabilized blood samples would experience in transit using a typical courier service (see rationale for design described at the beginning of the Results and Discussion). The samples underwent external changes in temperature between 21°C and 41°C over the course of three hours before storage at -80°C until the time of analysis (Fig. 2A). Of the 8 participants, stabilized blood samples from 7 of the donors were exposed to these high environmental temperature spikes. Samples 1 and 2a (from participant 2 who was sampled twice: 2a for control and 2b for temperature stress) were frozen after overnight incubation at room temperature to serve as controls.

**Figure 2.**
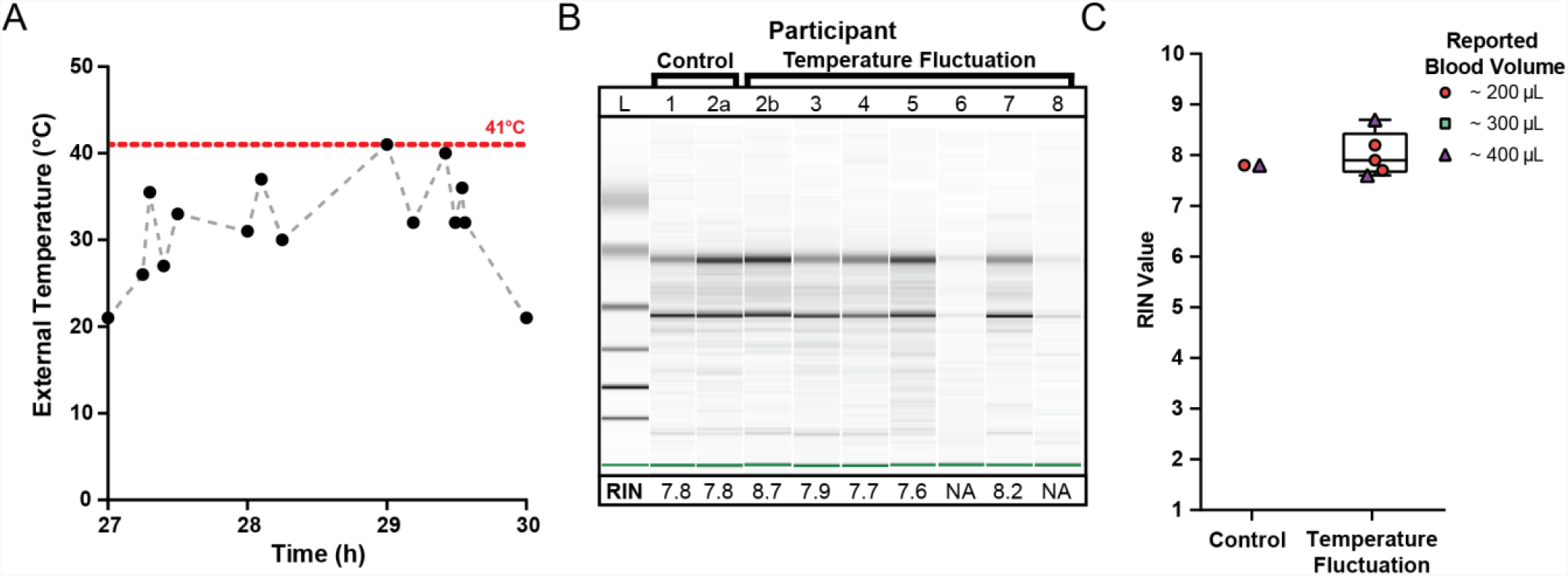
Quality of isolated RNA from stabilized *home*RNA samples exposed to high external temperature spikes in Doha, Qatar. A) External temperature fluctuation experienced by self-collected and stabilized blood samples after storage at ambient temperature (21°C) for 27 h. External temperature reached a maximum of 41°C. Black data points indicate the measured temperature; gray dashed lines are included to connect the points to guide the eye only (transitions between temperatures may not be linear). B) Digital gel image of RNA isolated from *home*RNA blood samples with corresponding RIN values. Samples 2a and 2b were from the same participant. Samples 1 and 2a did not undergo temperature fluctuations (they were frozen down at -80 C after overnight ambient temperature incubation to use as controls). C) Scorable RIN values for each blood sample not exposed (control) and exposed (temperature fluctuation) to high temperature spikes along with the reported approximate blood volume collected by each participant. Participants 6 and 8 yielded too low of a RNA concentration to be detected using the RNA 6000 Nano kit.

The quality of isolated RNA from all 9 blood samples (from 8 participants) was assessed to observe if the high external temperature spikes had adverse effects on the RNA stability in the *home*RNA platform. Of the 9 samples, 7 yielded scorable RIN values (Fig. 2C). The two that did not yield a RIN value (participants 6 and 8) both reported collected blood volumes of ∼100 μL. We note that blood volume does not necessarily correlate with RNA yield (see Figure S5) and that variability in blood volume could result from device-to-device variability in the Tasso blood collection system and inaccuracy in participant reporting of the volume of collected blood. These two samples had low RNA concentrations beneath the limit of detection of the RNA 6000 Nano Kit (<5 ng/μL), but distinct 18S and 28S rRNA bands are noted in the digital gel (Fig. 2B). The scorable RNA samples had RIN values between 7.6 - 8.7, demonstrating a sample quality suitable for downstream transcriptome (e.g., RNA-seq) analysis. Electropherograms obtained from the bioanalyzer can be found in the SI (Fig. S1). RNA concentrations for the first 50 μL elution were also obtained with the bioanalyzer (Fig. S2). For the 5 samples that were exposed to temperature fluctuations and yielded a scorable RIN value, the RNA concentrations from the first 50 μL elution ranged from 21 ng/μL to 40 ng/μL (1050-2000 ng total cellular RNA in the first 50 μL elution). The two samples that yielded a concentration less than the qualitative range (5 ng/μL) of the Nano 6000 kit (participant 6 and 8) both measured a concentration of 4 ng/μL (200 ng total cellular RNA in the first 50 μL elution). Therefore, all samples, including the low concentration samples (participant 6 and 8), likely had at least 200 ng total yield of RNA in the first elution. In general, 500 ng is a comfortable minimal cutoff value for large-scale transcriptomics analyses such as standard RNA-Seq (with many facilities accepting lower amounts of RNA), and 100 ng is a comfortable minimal cutoff value for expression analyses of a small panel of targeted genes. For RNA-Seq, we note that different cutoff values exist (for example as low as 10-30 ng)depending on the method and whether cDNA amplification is performed.

The successful isolation and high quality of RNA from blood samples collected, stabilized, and analyzed in Doha suggests that the *home*RNA platform can be used for at-home clinical studies in Doha or places similar to Doha, a hot climate where air conditioning is commonplace in buildings and vehicles and fast courier services are available. In this setting, samples experience exposure to short term, high temperature fluctuations in external environmental conditions. Perhaps some researchers would assume that rapid temperature spikes would not affect the quality of RNA*later*™-stabilized RNA, but we felt that it was necessary to conduct this field testing before using *home*RNA for clinical research in Doha.

One important note regarding the experimental setup for this small pilot study is that the reported temperatures (Fig. 2A, Table S1) are that of the local external environment (i.e., temperature inside the car, ambient outdoor temperature, etc.) rather than the temperature of the blood sample itself. It is possible that even though the external temperatures fluctuated markedly over the course of three hours with intervals of rapid but short increases or decreases in ambient temperatures, the blood sample itself may not have fully equilibrated to these observed external temperatures. Further experimentation, such as in-lab controlled temperature stress experiments, could further determine how spikes in external temperature affect the temperature of the blood sample itself.

### home*RNA for high temperature seasons: a pilot study in Western and South Central USA*

To further assess the robustness of the *home*RNA kit in high temperature settings, 11 participants in various regions throughout the Western and South Central USA collected and stabilized their own blood and shipped it back to our lab for analysis. These samples were collected during the summer of 2021, which involved unusually high temperatures in some locations, and 2 of these samples were collected during the June 2021 heatwave in the Pacific Northwest region. Samples were collected at indoor ambient temperatures ranging between 21°C and 27°C (Table 1, Fig. 3Bi). Typically, stabilized blood samples were shipped back the same day or the following day with overnight shipping. The logistics of the study were such that participants left their package in a designated pickup area; all participants chose to have their samples picked up from outside their homes; therefore, all the stabilized blood samples were also exposed to the outdoor ambient temperatures following indoor collection and stabilization (See Table 1 for pickup location). The duration of post-collection specimen storage prior to transit back to the lab (turnover duration) can be highly variable; this ranges from immediate transit (direct package handover a courier service driver) to overnight storage, in which these specimens can experience prolonged exposure to high ambient temperatures. This turnover duration is an important variable in temperature-dependent specimen stability and our data reflects the cumulative effect of variations in turnover duration experienced during real-world transits.

**Figure 3.**
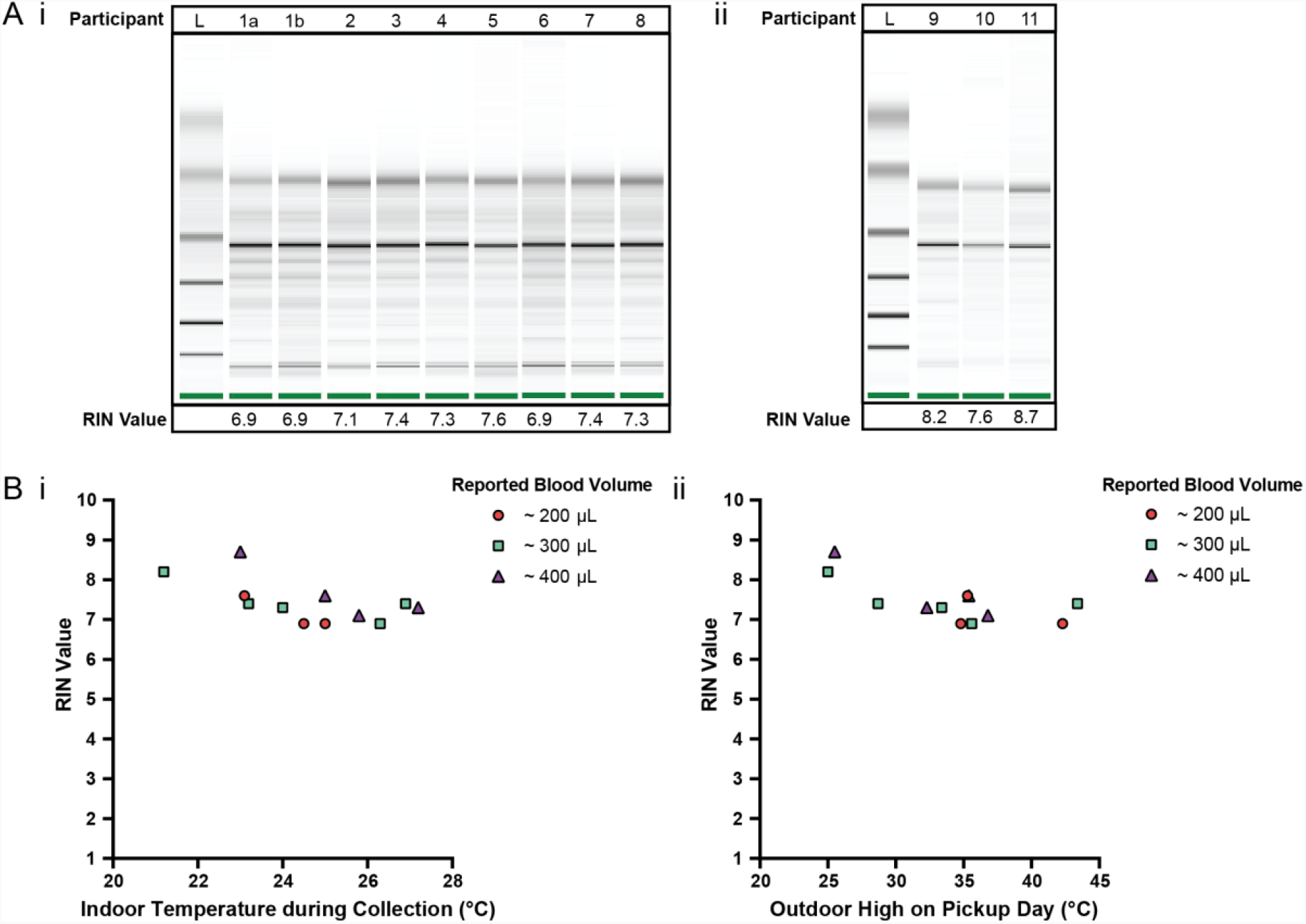
Quality of isolated RNA from stabilized *home*RNA samples collected during the summer in the Western and South Central USA. A) Digital gel image of RNA isolated from *home*RNA stabilized blood samples with corresponding RIN values, with i) samples diluted 1:5 with water and run with a RNA 6000 Pico kit and ii) samples run with a RNA 6000 Nano kit without dilution. Participant 1 contributed two samples, 1a and 1b, that were collected and stabilized on different days and at different temperatures. B) i) RIN values from blood collected and stabilized at different indoor ambient temperatures during summer and ii) RIN values according to the outdoor temperature high reported on the day of collection. Each participant was asked to report the approximate blood volume collected before stabilization based on Fig. S6 in the collection survey.

According to the day that each sample was collected and stabilized, the maximum outdoor temperature (“daily high”) reported for the corresponding location of each participant (based on local weather reports found at Weather Underground) is included in Table 1. These outdoor temperatures spanned between 25°C and 44°C (Table 1, Fig. 3Bii). We note that we do not know if the sample was outside during the daily high and further note that it is possible that the package was exposed to temperatures higher than the daily high, such as if it was placed in direct sunlight or if the temperature inside the pickup vehicle exceeded the daily high. Thus, the daily high is simply provided as a reference. In addition to the above, as noted in the Doha pilot study, the temperature of the blood sample may be different from the temperature of the external environment.

The quality of isolated RNA from 12 blood samples (from 11 participants) was assessed to observe how various temperature exposures (indoor, outdoor, shipping conditions, etc.) affected *home*RNA stability in a field test. 75% of the samples (9 out of 12 samples) yielded a RIN ≥7.0 (7.1-8.7) and the remaining 3 samples yielded a RIN of 6.9 (Fig. 3A). The RIN values demonstrate sufficient RNA quality for downstream transcriptomic analysis.

To observe how temperature at the time of collection and shipment affected the RNA integrity, RIN values were plotted against the respective indoor temperature at the time of collection and stabilization (Fig. 3Bi) along with the local weather reports maximum outdoor temperature (Fig. 3Bii). Indoor temperatures reported for the 12 samples range from 21.2-27.2°C. Indoor temperature at the time of collection (at least across the temperature range and small sample size used in this study) does not have a clear influence on the RIN value. Notably, there is overlap between the indoor temperatures reported here (Fig. 3B) and in our initial *home*RNA feasibility study ^23^. In our previous pilot study conducted in 2020 ^23^, the majority of the samples were collected with an ambient indoor temperature between 18-25°C (n = 43/53 analyzed returned samples where participants completed their surveys and provided indoor temperature data), with the other 9 samples ranging from 26-28°C, with 5 at 26°C, 1 at 27°C and 3 at 28°C. The results from the present study conducted in 2021 with respect to indoor temperature are plotted together with the results from our 2020 pilot study ^23^ with regards to RIN (Fig. S5Ai) and total RNA yield (Fig. S5Bi), in the supporting information. There is also some overlap in the range of outdoor highs on pickup day in this study and our previous study conducted in 2020, but the samples collected in the 2021 study described here in the Western and South Central USA were picked up on days with a warmer outdoor high, on average. For the 2020 pilot study, the outdoor highs on pickup day ranged from 9°C-34.8°C, with a median outdoor high of 21.4°C, and only 7/60 total samples (collected from 47 total participants) had an outdoor high on the pickup day >30°C ^23^. In contrast, the median and range of the outdoor high on pickup day was higher for the present 2021 study, ranging between 25-43.4°C, with a median of 35.1°C. The RIN (Fig. S5Aii) and total RNA yield (Fig. S5Bii) from the 2020 and 2021 studies are plotted together with respect to outdoor high on pickup day in Fig. S5. All but one of the samples where a RIN was obtained in our 2020 pilot study yielded a RIN ≥7.0; the one sample with a RIN <7 was evaluated to a RIN of 6.8^23^. With regards to the relationship between the outdoor high and RIN in the present study, the three samples with the lowest RIN (RIN of 6.9) correspond to outdoor daily high temperatures ≥34°C; however, four other samples corresponding to outdoor daily high temperatures ≥35°C returned RIN values ≥7.0.

Although RIN values were indicated as the main performance metric in this study, we acknowledge that other performance metrics (e.g., total yield and RNA concentration) may be used concurrently to guide sample selection for downstream analysis. In longitudinal studies with multiple collection time points, extremely low yields either due to low blood volume or low circulating WBCs can lead to exclusion of specific time points in the longitudinal data series. These missing-at-random (MAR) samples can be problematic in longitudinal studies as they create unbalanced data and biased estimates ^30^. However, additional statistical approaches such as time-course gene set analysis (TcGSA) can be used to account for such heterogeneity in the dataset ^30^.

In clinical studies where sampling is done in high temperature climates and samples are susceptible to variable levels of temperature-induced degradation, it is crucial that initial analysis of the genes of interest is performed to determine how their transcript levels change with respect to post-stabilization temperature stress as degradation may not occur uniformly within the sample ^29^. Prior to complete sample stabilization, multiple RNA decay pathways may influence rate of decay for distinct transcript sets ^31^. In such studies, alternative normalization strategies (e.g., using linear model framework to control for effects of RIN values) may be used to enable identification of biological meaningful expression signatures ^29^.

## CONCLUSION

The ability to remotely collect and stabilize blood will expand future applications in the field of personalized medicine, particularly for analyzing blood transcriptomes that can give insight to disease progression/therapy response, immune response, and biomarker discovery. However, in order to establish the capability to do this, it is important to examine how exposure to various temperature conditions affects RNA stability. In this work, we conducted two small pilot studies to test the robustness of our *home*RNA at-home self-collection and stabilization blood kit to successfully preserve RNA integrity in high ambient temperature environments (>30°C) in two different study designs. These two pilot studies can inform clinical studies that could be run in places similar to Doha, Qatar (where samples are only exposed to short temperature spikes in transit) and the Western and South Central USA (where samples are in transit overnight given the regional nature of the study). We demonstrated that the majority of these blood samples yielded sufficient RIN (≥7.0), which suggests the usability of these samples for downstream transcriptome analyses. Important limitations of the present work include relatively small sample sizes, although we note that this work follows on our larger study with 47 participants across the USA ^23^. Further, we have several ongoing clinical studies in the USA, with >1000 samples collected over the past year; these studies will eventually add to the literature on the robustness of *home*RNA across the USA through different seasons. It is also important to study the performance of *home*RNA in tropical climates where samples will undergo longer transit times and longer periods of sustained high temperatures than those expected in Doha; to this end, we have ongoing studies utilizing *home*RNA in urban and rural Thailand. As *home*RNA is applied to diagnosis and monitoring of specific diseases, it will be important to analyze the expression of genes of interest that could be affected by elevated temperatures (i.e., transcriptional levels of inflammatory genes or disease-specific genes) using controlled in-lab experiments prior to personalized medicine applications. To our knowledge, this is the first study to establish use of *home*RNA in sampling regions experiencing higher ambient temperatures, representing an initial validation study to extend the use of *home*RNA to warm environments.

## ACKNOWLEDGEMENTS

This publication was supported by the David and Lucile Packard Foundation, an Alfred P. Sloan Research Fellowship, the University of Washington, a STEP grant from UW CoMotion, and the National Institutes of Health (NIH) through the University of Washington EDGE Center of the National Institute of Environmental Health Sciences (P30ES007033) and the National Center for Advancing Translational Sciences award number UL1 TR002319. The content is solely the responsibility of the authors and does not necessarily represent the official views of the National Institutes of Health. We would like to acknowledge Proudrawee Suthakorn for the figure drawings in Fig. 1. We would also like to acknowledge Grant Hassan for assisting with study logistics for collecting the Western and South Central USA samples, as well as Grant Hassan, Hannah Lea, and Ashley Dostie for their help in making and shipping the *home*RNA kits to Qatar. We would also like to thank Lochlan Hickok, Paul Miller, Annalyn Randal, Madeleine Eakmen, Mike Zimmerman and Angela Mullen for helping facilitate shipping, logistics, and procurement specific to our remote sampling study.

