## Supplementary Material for "At-home blood collection and stabilization in high temperature climates using *home*RNA"

### Table of Contents:

Supplemental Tables and Figures Pg. S2 – S7
References Pg. S7
Instructions for Use Pg. S8 – S9

**SUPPLEMENTAL TABLES AND FIGURES:**

**Table S1: Participant and sampling information for the Western and South Central USA pilot study**

| Time (h) | Event / Procedure | External Temperature (℃) | Location |
|---|---|---|---|
| 0 | Overnight hold at room temperature | 21 | Lab, Research |
| 27 | Start in lab | 21 | Lab, Research |
| 27.25 | Transport outside to car | 26 | Outside |
| 27.3 | In car - A/C started | 35.5 | Indoor parking |
| 27.4 | In car - A/C on | 27 | Indoor parking |
| 27.5 | In car - driving with some sun exposure | 33 | Driving |
| 28 | In car - A/C stopped | 31 | Outdoor parking - moderate sun exposure |
| 28.1 | In car - A/C restarted | 37 | Outdoor parking - moderate sun exposure |
| 28.25 | Car stopped - A/C off | 30 | Outdoor parking - covered |
| 29 | Car restarted - A/C on | 41 | Outdoor parking - covered |
| 29.19 | Car stopped - A/C off | 32 | Home parking - covered |
| 29.42 | Car restarted - A/C on | 40 | Home parking - covered |
| 29.49 | Car stopped - A/C off - Walking to Lab | 32 | Indoor parking |
| 29.54 | Enter building | 36 | Research |
| 29.56 | Office | 32 | Office, Research |
| 30 | Transferring blood from stabilizer tube to store | 21 | Lab, Research |
| 30.3 | Storing sample | -80 | Lab, Research |

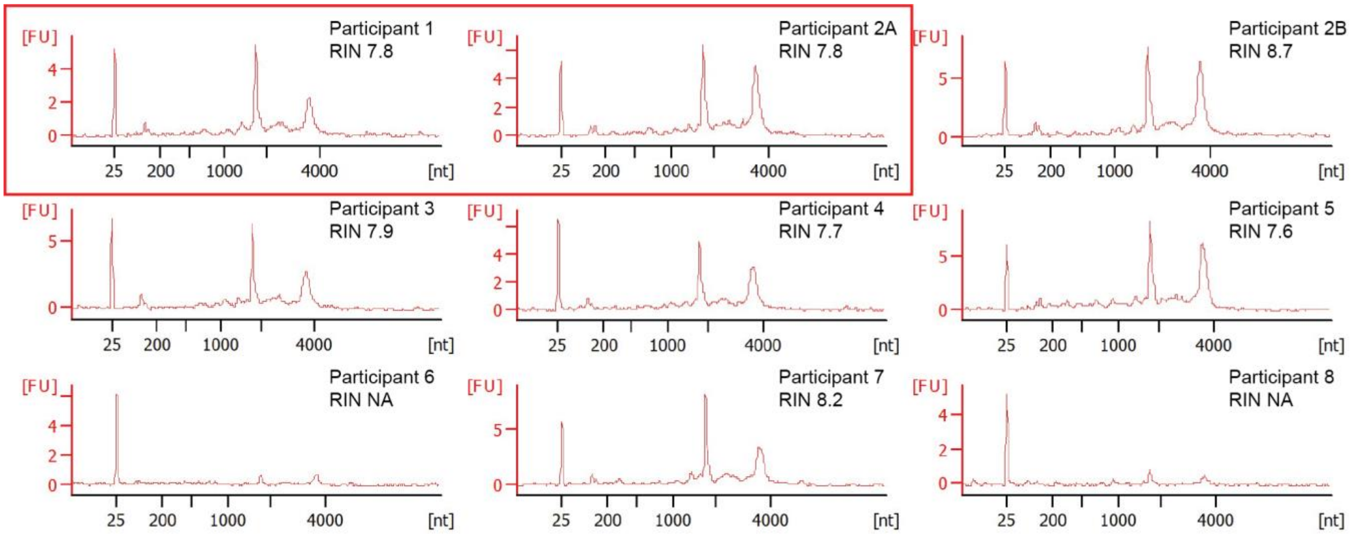


**Figure S1. Electropherograms of isolated RNA from stabilized *home*RNA samples in Doha, Qatar.** Electropherograms were obtained as raw data using a RNA 6000 Nano Kit on an Agilent 2100 bioanalyzer, which uses fluorescence to detect the marker (~25 nt), 18S rRNA fragments (~1900 nt), and 28S rRNA fragments (~3900 nt). The RIN algorithm then uses these peaks to assign a RIN value to the corresponding sample. The electropherograms are used to generate a gel image, as seen in Fig. 2B. Samples 1 and 2A in the red box correspond to control samples.

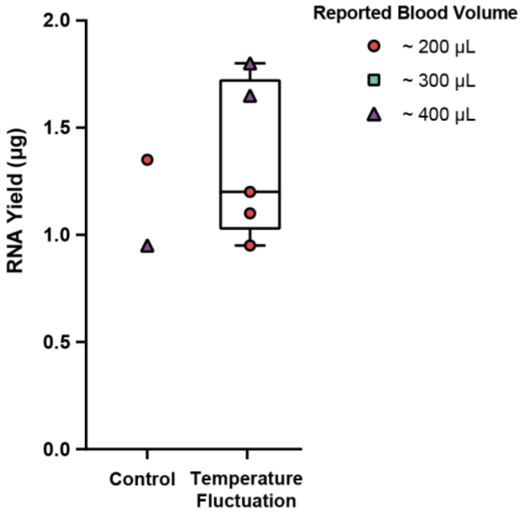


**Figure S2. Yield of isolated RNA from stabilized *home*RNA samples in Doha, Qatar.** Reported yields were obtained from bioanalyzer measurements by taking the reported concentration from the first 50 µL elution. For the two samples (from participant 6 and 8) that did not have scorable RIN values, yields were not included on the graph since their concentrations were below the lower limit of the qualitative range of the RNA 6000 Nano kit (5 ng/µL). However, for reference, the bioanalyzer reported 4 ng/µL for participants 6 and 8. While concentrations from the Agilent 2100 Bioanalyzer are not typically used for obtaining yield, a threshold value of 200-500 ng total yield is typically needed for downstream analysis such as RNA sequencing. Therefore, an exact value is less important and the bioanalyzer can be used to obtain approximate RNA concentration ranges.

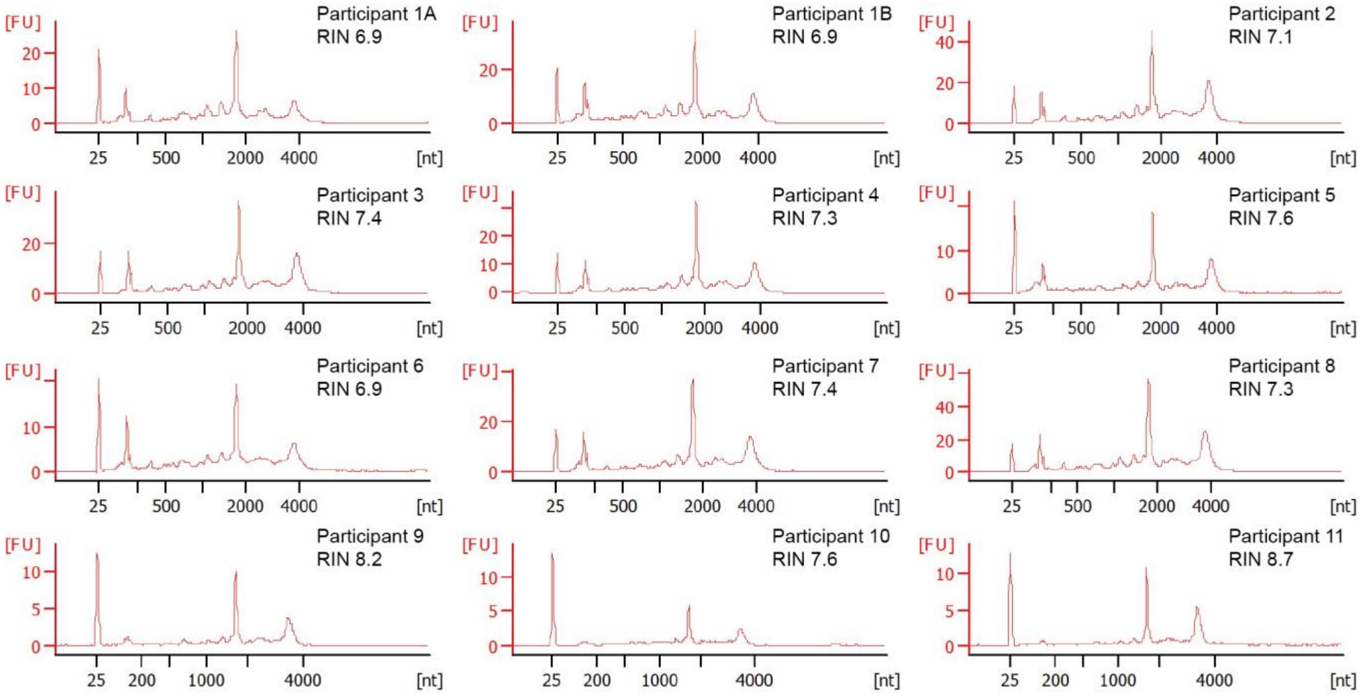


**Figure S3. Electropherograms of isolated RNA from stabilized *home*RNA samples in the Western and South Central USA.** Electropherograms were obtained as raw data using a RNA 6000 Pico Kit for samples 1-8 and a RNA 6000 Nano Kit for samples 9-11 on an Agilent 2100 bioanalyzer. For samples 1-8, the RNA 6000 Pico kit uses fluorescence to detect the marker (~25 nt), 5S rRNA fragments (~150 nt), 18S rRNA fragments (~1900 nt), and 28S rRNA fragments (~3900 nt). The RNA 6000 Nano kit detects the same peaks with the exception of the 5S rRNA fragment peaks. The RIN algorithm then uses these peaks to assign a RIN value to the corresponding sample. The electropherograms are used to generate a gel image, as seen in Fig. 3A.

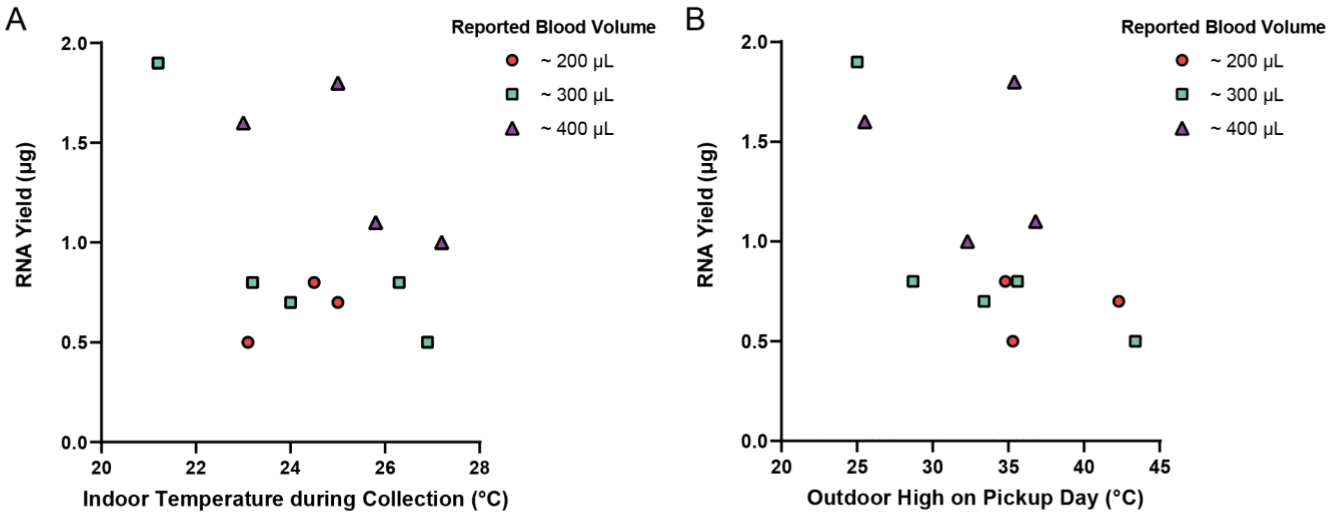


**Figure S4. Total yield of isolated RNA from stabilized *home*RNA samples collected, stabilized, and shipped at high temperatures in Western and South Central USA.** Reported total yields were obtained by measuring both 50 µL elutions from the RNA isolation protocol (see Methods and Materials) on Cytation 5 Multi-Mode Reader and adding them together to obtain a total yield. Each total yield is reported with the corresponding A) indoor ambient temperature at the time of collection and B) reported outdoor temperature high on the pickup day.

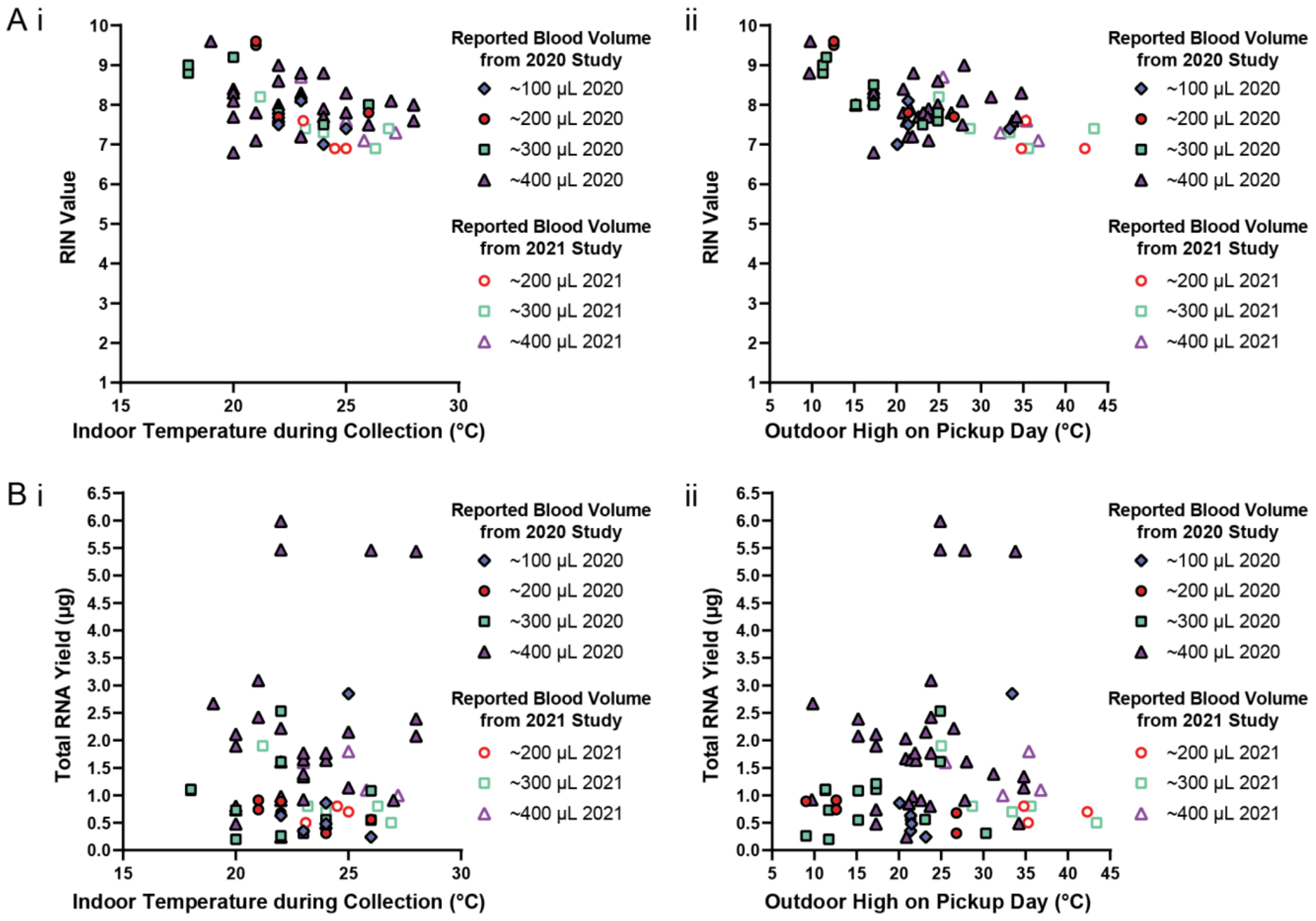


**Figure S5. RNA quality and quantity versus temperature, combining data from the present study (summer 2021) from our previous pilot study (summer 2020, Haack, Lim *et al.* 2021).**[1] A) RIN values versus i) indoor temperature and ii) outdoor high on pickup day. A) Total RNA yield versus i) indoor temperature and ii) outdoor high on pickup day.

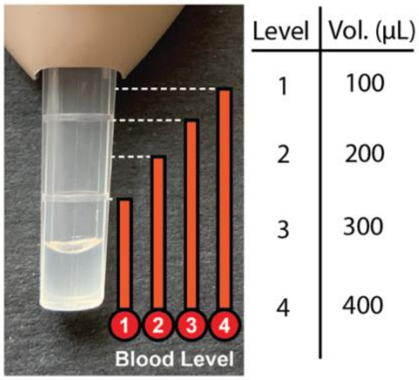


**Figure S6. Reported blood level volumes.** Study participants for both the Qatar and Western and South Central USA groups were asked to report the approximate volume with each blood collection with the Tasso-SST™ based on this picture provided in the survey that they completed with sampling. Figure was reprinted with permission from Haack, Lim *et al.* *home*RNA: A Self-Sampling Kit for the Collection of Peripheral Blood and Stabilization of RNA**.** *Anal. Chem.* 2021, 93, 39, 13196–13203. Copyright 2021 American Chemical Society.[1]

#### INSTRUCTIONS FOR USE:

The IFU for the self-collection and stabilization of blood using *home*RNA is shown below (Pg. S8 – S9). Figure was reprinted with permission and adapted from Haack, Lim *et al.* *home*RNA: A Self-Sampling Kit for the Collection of Peripheral Blood and Stabilization of RNA**.** *Anal. Chem.* 2021, 93, 39, 13196–13203. Copyright 2021 American Chemical Society.[1]

#### PREPARE DEVICES AND APPLICATION SITE

**FIRST TIME USERS**

Please watch the instructional video provided at

https://www.youtube.com/watch?v=iV3GZ8SmmuM

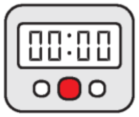


**1. Wash hands and get a timer.**

You will use a timer in Steps 2 and 10.

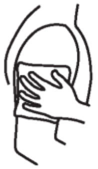

**2. Apply hot pack to upper arm for 2 minutes.**

Warming helps your blood flow better.

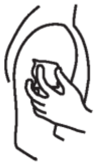

**3. Clean arm with alcohol wipe.**

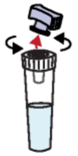

**4. Open the stabilizer tube by twisting off the purple cap.**

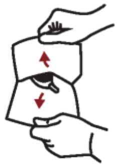

**5. Open Tasso pouch by pulling apart white and clear layers.**

Discard cap in pouch.

#### COLLECT BLOOD USING TASSO-SST

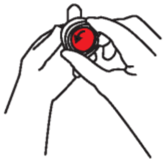

**6. Remove clear plastic cover over the red button.**

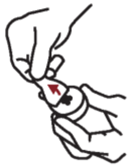

**7. Peel paper tab behind the red button.**

Keep the tube pointing down.

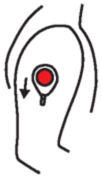

**8. Stick device to shoulder.**

Do not remove once it is on.

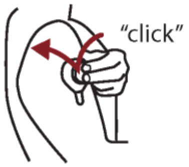


**9. Press button quickly and firmly until it can't go any farther. Wait 2 seconds then let go.**

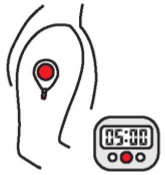


**10. Start a 5 minute timer.**

Keep arm at your side. You wont see blood right away. It can take up to a minute for blood to flow.

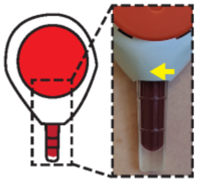

**11. After 5 minutes or when the tube fills, *whichever comes first*, peel off the device.**

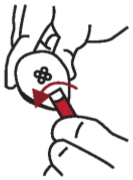

**12. Remove tube by firmly twisting a quarter turn and pulling down.**

This may take a bit of finger strength.

#### MIX AND PACKAGE

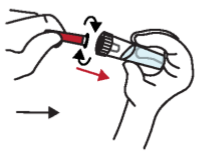

**13. Bring together the blood tube and stabilizer tube and screw these together tightly.**

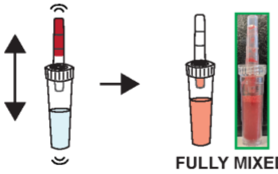


**14. With the stabilizer tube on the bottom, shake hard up and down to mix. Stop when mixed.**

Some fluid may remain in blood tube. When mixed, the color is the same. See insert for details.

**DO NOT DISCONNECT TUBES.**

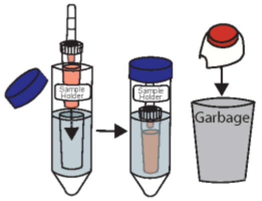


**15. Place sample in sample holder. Throw away used Tasso device.**

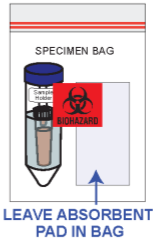


**16. Place blood sample in the specimen bag. Wash your hands with soap and water**

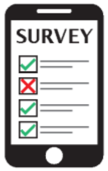


**17. Fill out the online Collection Survey that was emailed to you.**

You will need the Unique Code on the box.

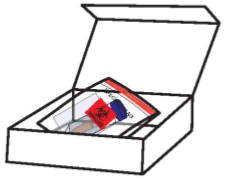

**18. After filling out the survey, place your sample back in the box, and return using the provided shipping return bag.**

For research use only

### INSTRUCTIONS FOR USE

#### Home Blood Collection Kit

If you need assistance please contact us at or call [redacted]

**Thank you for participating in our study!**

#### KIT CONTENTS

Make sure your kit contains all components listed below

**1) Tasso-SST device**

**2) Stabilizer tube**

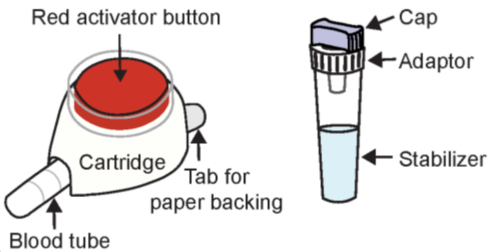


**3) Alcohol wipes**

**4) Hot pack**

**5) Bandage**

**6) Specimen bag**

**7) Sample Holder**

**8) Mail return bag**

#### INTENDED USE

TASSO-SST is a single use blood collection device that is intended for the self collection of capillary blood from the upper arm of adults (18 years or older). The stabilizer tube contains liquid that is intended for stabilizing the collected blood. The home blood sampling kit is for academic research use.

#### STORAGE

Store at 15 - 30ºC (60 - 80ºF) in a dry place.

#### WARNINGS

- The TASSO-SST is a sterile device. Do not open until use.
- The TASSO-SST device contains sharps. Handle with care.
- For external use only.
- Keep out of reach of children.
- Use while seated as fainting may occur during blood sampling procedure.
- Do not activate the red button until device is firmly on skin.
- Wipe application site with alcohol wipe to reduce infection risk.
- Always use a new unopened pouch of Tasso-SST. Do not re-use.

#### ONLINE SYMPTOM AND COLLECTION SURVEY

Please check your e-mail or text message based on the preference you has listed for a link to fill out the daily use online survey. If you cannot find the link, e-mail us at [redacted]
